# Parasite epigenetic memory and blood barriers dictate host transcriptional responses during generalist host-shifts

**DOI:** 10.64898/2026.08.27.747310

**Authors:** L García-Longoria, J Aželytė, V Palinauskas, O Hellgren

## Abstract

The evolutionary success of generalist parasites is often attributed to their capacity to rapidly navigate divergent host environments through transcriptional plasticity. While host-parasite dynamics are frequently studied in avian models, the immunogenic impact of the heterologous blood matrix, a critical variable in cross-species inoculation experiments, is rarely accounted for. In this study, we investigated the host-transcriptomic landscape of domestic canaries (*Serinus canaria*) infected with the avian malaria parasite *Plasmodium homocircumflexum* (lineage COLL4), employing a crosswise infection design to differentiate between homologous and heterologous donor sources. By implementing a factorial experimental framework, we successfully isolated the transcriptional noise induced by the heterologous blood matrix per se, revealing that mismatched transfusions trigger significant, non-specific innate immune activation independently of parasite presence. Upon correcting for this background effect, we observed distinct transcriptional trajectories: while adapted (homologous) infections induced a metabolic catalytic overload driven by key kinase hubs (e.g., AKT1, CDK6), heterologous infections were characterized by a shift toward structural and ribosomal regulation. These divergence patterns in the host, combined with the strain-specific transcription of the parasite, suggest that early infection phases are heavily constrained by recent host-switching events. Our results demonstrate that this epigenetic memory acts as a fundamental determinant of virulence, providing a new systems-based framework for understanding how pathogen history and host-donor compatibility reshape infection dynamics and host molecular outcomes during the colonization of novel ecological frontiers.

## Introduction

The evolutionary success of generalist parasites hinges on their ability to rapidly adapt to divergent immunological and metabolic landscapes following a host switch (Prati et al. 2022). Rather than relying on immediate genomic alterations, this adaptive flexibility is frequently driven by transcriptional variations and complex epigenetic mechanisms that allow the pathogen to navigate a novel host’s immune response (Adejoh et al. 2023). For instance, the transcriptome of specialized blood parasites displays distinct host-specific gene expression, illustrating how plastic phenotypes facilitate survival in novel environments (Videvall et al. 2017). Increasingly, advanced molecular protocols for detecting and characterizing these lineages in the field are providing an ever-growing picture of their prevalence, distribution, host range, and diversity hotspots worldwide (Musa et al. 2024; Pacheco et al. 2024; Emmenegger et al. 2026). Consequently, these blood-borne organisms have re-emerged as unparalleled experimental models for addressing evolutionary ecology under varying selective pressures (Rivero and Gandon 2018), including critical life-history adjustments such as sex determination (Paul et al. 2000), fertility insurance shaping sex ratios (West et al. 2002), and the fine-tuning of transmission timing to match vector fluctuations (Pigeault et al. 2018). Deciphering how these hidden regulatory layers orchestrate such rapid multi-host lifestyles promises to reshape our entire understanding of how pathogens conquer new ecological frontiers.

To unravel the complexities of these host-parasite dynamics, certain avian haemosporidian systems serve as a paradigm for coevolutionary research (Valkiūnas 2005). This globally distributed group of blood pathogens has not only restructured our understanding of host specificity, virulence, and parasite dispersal (Martinsen et al. 2008; Svenson-Coelho et al. 2013; Ellis et al. 2019), but it also allows researchers to time the evolutionary radiation of these diverse lineages across the globe (Pacheco et al. 2018; Pacheco and Escalante 2023). Unlike traditional rodent models or heavily treated human populations, wild avian systems enable the study of natural selection without the confounding interference of anthropogenic measures like vaccines or vector control (Ayala et al. 2016; Kyriazis et al. 2025). Field investigations are further enhanced by high parasite prevalence, ease of access, and the ability to mark and recapture hosts within a deeply established ornithological framework (Asghar et al. 2011; Renner et al. 2026). This unique animal model allows researchers to manipulate host physiology while minimizing parasite genetic variation by inoculating different host species with the exact same lineage (Palinauskas et al. 2008; Videvall et al. 2017). Consequently, by accounting for confounding variables such as age, dietary intake, and prior exposure (McGraw and Ardia 2003; Owen-Ashley et al. 2004; Cornet et al. 2014; Ellis et al. 2015), avian models might demonstrate how identical pathogen genotypes trigger drastically different virulence profiles across distinct host species, moving past the taxonomic limits of traditional molecular frameworks (Outlaw and Ricklefs 2014).

When evaluating vector-borne infections via experimental inoculations, however, the immunogenic effect of the blood matrix itself is frequently overlooked. While birds lack direct homologs of the human ABO blood group system (Yamamoto 2017), historical frameworks in chickens (*Gallus gallus*) (Dietert and Dietert 1992; Taylor et al. 2016) captured only a fraction of this complexity. Recent advances reveal that modern birds (Neoaves) express sophisticated counterparts to the P1PK blood group system on their red blood cells (Bereznicka et al. 2026). Because these glycosphingolipids vary widely across species and environments, introducing a heterologous blood matrix can trigger confounding immune artifacts (Bereznicka et al. 2026). Exposure to mismatched donor blood induces anti-P1 or anti-Pk responses, while environmental exposure to cross-reactive microbiota further primes hosts to generate antibodies against foreign erythrocyte structures (Mäkivuokko et al. 2012; Cabezas-Cruz et al. 2017; Wu et al. 2018). Consequently, the “blood matrix per se” acts as a critical, unquantified biological variable in dual transcriptomics; failing to account for donor-recipient compatibility risks masking true pathogen-specific host responses behind background transfusion-like immune reactions.

Crucially, this baseline host reactivity does not occur in a vacuum; it directly collides with the parasite’s own adaptive machinery during the host-switching window. Beyond immediate tissue environments, a parasite’s survival hinges on its capacity to navigate varying immune responses, body temperatures, and nutrient availabilities during a host shift (Agosta et al. 2010; Gupta et al. 2020; Prati et al. 2022). To overcome these challenges across distinct species (Palinauskas et al. 2020; Galinski 2022) and geographic regions (Clark et al. 2020; Fecchio et al. 2021), vector-transmitted blood pathogens rely heavily on transcriptional variation (Mandala et al. 2021; Turnbull et al. 2022; Matos et al. 2023). In this sense, longitudinal data reveals that early-stage gene expression in the parasites remains strictly donor-dependent before gradually shifting to a recipient-aligned profile (García-Longoria et al. 2025). This lag suggests that clonal pathogen “offsprings” after asexual reproduction retain an epigenetic “memory” or imprinting from their previous host—mediated by mechanisms like DNA methylation (Serrano-Durán et al. 2022) and histone modifications (Connacher et al. 2022; Hollin et al. 2023) – where natural selection acts directly on regulatory variation rather than underlying haplotypes (Nourmohammad et al. 2017; Cope et al. 2025). While vector-stage transitions further refine these dynamics (Yu et al. 2022), initial donor-driven synchronization proves that a parasite’s historical journey shapes the outcome of an infection just as profoundly as its genome, offering a fresh framework for unraveling the hidden constraints behind pathogen specialization and host-switching kinetics.

Thus, the overarching goal of this study is to characterize the host transcriptomic landscape during a protozoan infection and determine how the origin of the pathogen population shapes the host’s molecular response. To achieve this, we have defined four specific objectives: (i) to characterize the temporal transcriptomic response of the domestic canary (*Serinus canaria*) across the early, peak, and resolution stages of infection; (ii) to evaluate how the previously host source of the parasite inoculum influences host gene expression by comparing infections derived from homologous versus heterologous donors; (iii) to quantify the independent immunogenic effect of the blood matrix, using uninfected controls to isolate the transcriptional “noise” generated by heterologous blood injection *per se*; and (iv) to perform a dual host-parasite integrative analysis linking host molecular trajectories with the pathogen’s transcriptional state. By decoupling these confounding variables, this research aims to uncover the fundamental regulatory boundaries that define vector-borne host-switching events.

## 2. Materials and Methods

### 2.1 Experimental Animals and Ethical Approval

The study was conducted at the State Scientific Research Institute Nature Research Centre in Vilnius, Lithuania. Experimental subjects included juvenile domestic canaries (*Serinus canaria*) and Eurasian siskins (*Spinus spinus*), both of which were confirmed to be free of blood parasites via microscopy and nested PCR prior to the start of the experiment. All experimental protocols followed European and Lithuanian regulations for animal research and were approved by the State Food and Veterinary Service of Lithuania (No. 2018/05/03-G2-84).

### 2.2 Experimental animals and parasite acquisition

We used the avian malaria parasite *Plasmodium homocircumflexum* (lineage COLL4, GenBank KC884250), originally isolated from a red-backed shrike (*Lanius collurio*). The strain was cryopreserved following Dimitrov et al. (2015) and later used for experiments. To investigate the host response to adapted versus heterologous infection, a crosswise infection experiment was implemented (see more information in García-Longoria et al. 2025). Host canaries were randomly assigned to four treatment groups (4 birds per group) (Figure 1): (i) Ec: Canaries infected with blood from an infected canary donor. (ii) Es: Canaries infected with blood from an infected siskin donor. (iii) Kc: Canaries inoculated with clean canary blood. (iv) Ks: Canaries inoculated with clean siskin blood. Sampling was performed at day 0 for control groups, and at days 8, 12, and 16 post-inoculations for the experimental groups. Birds were housed individually in cages, allowing social interaction, in a vector-free room with a stable 21±1 °C temperature and a natural light/dark cycle. Food and water were provided *ad libitum*.

**Figure 1.**
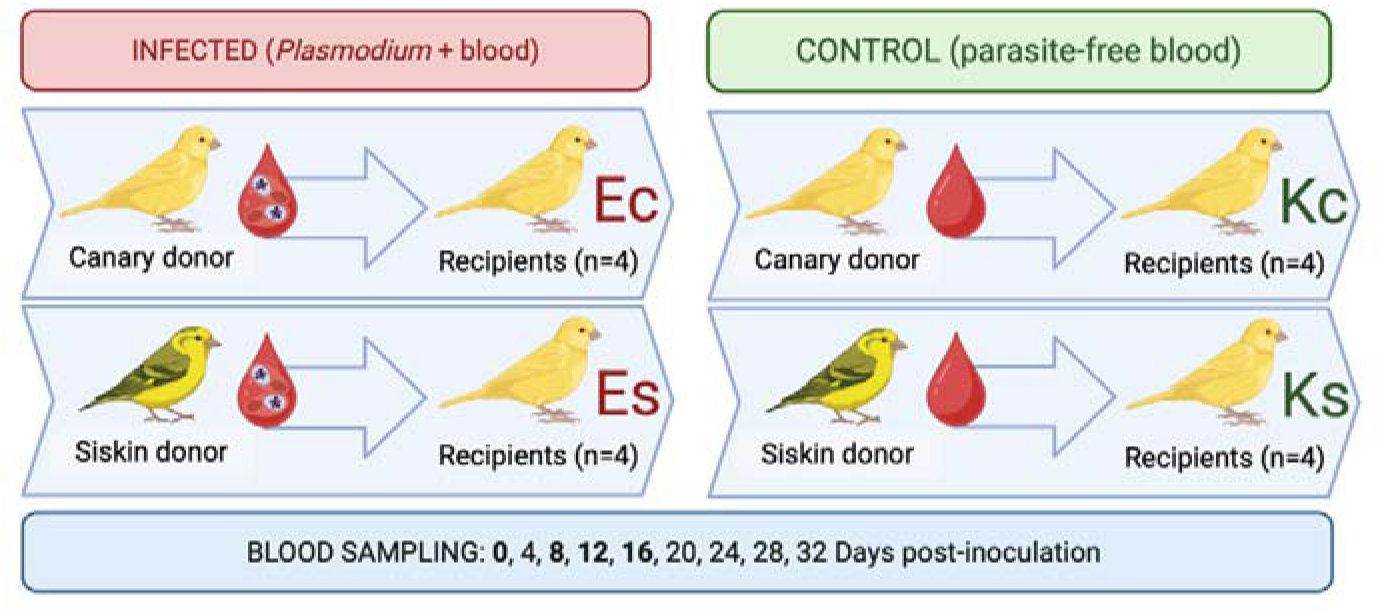
Experimental design of the inoculation study. Schematic representation of the donor and recipient groups. Experimental groups (Ec and Es) were inoculated with *Plasmodium homocircumflexum* (left). Control groups (Kc and Ks) served as non-infected references for each donor species counterpart (right). Blood sampling was performed every 4 days (on days 0, 4, 8, 12, 16, 20, 24, 28, and 32 post-inoculation) to monitor the development of parasitemia. For the transcriptomic analysis, a specific subset of these samples was selected, corresponding to days 0 (only in control), 8, 12 and 16 (control and experimental).

### 2.3 Experimental procedures

The *P. homocircumflexum* strain COLL4 maintained since its original isolation via serial passage. To initiate infections, cryop reserved blood was introduced into a noninfected canary and siskin, which then served as donors: canary-derived parasites for canaries and siskin-derived parasites for canaries (Figure 1). Each bird received 100 μL of a freshly prepared suspension containing infected blood, 3.7% sodium citrate, and 0.9% saline (4:1:5 ratio), with an erythrocytic meront intensity of 0.4% and an estimated dose of 6×10^5^ meronts. The mixture was injected into the pectoral muscle following Iezhova et al. (2005). Birds were monitored for 32 dpi, with blood samples collected every 4 days for microscopy and PCR. Additional samples for RNA sequencing were taken at 8, 12, and 16 dpi.

### 2.4 Sampling, microscopy, and molecular analyses

Blood samples were obtained by puncturing the brachial vein. A small drop of blood from each bird was used to prepare two blood smears, while approximately 30 μl was stored in SET buffer (0.05 M Tris, 0.015 M NaCl, 0.001 EDTA, pH 8.0) for molecular analysis (Hellgren et al. 2004). Samples for RNA were immediately preserved in TRIzol LS Reagent (Invitrogen, Carlsbad, CA) and stored at –80 °C. Blood samples were kept at −20 °C until processed. The blood smears were air-dried, fixed with absolute methanol, and stained with Giemsa (Valkiu nas et al. 2008). The slides were examined under an Olympus BX51 light microscope equipped with an Olympus DP12 (Olympus, Shinjuku City, Japan). Approximately 100 fields were reviewed at a magnification of 1000×. Parasitemia was quantified by counting the number of parasites per 1,000 erythrocytes, or per 10,000 erythrocytes in cases of low infection intensity, following the protocol recommended by Godfrey et al. (1987). Detailed procedures for preparing, staining, and examining blood smears were outlined by Valkiu nas et al. (2008).

DNA extraction was performed using the ammonium acetate protocol (Sambrook et al. 1989). A nested PCR protocol was employed for genetic analysis (Waldenström et al. 2004). Amplification products (1.5 μl) were resolved on a 2% agarose gel. Sequencing followed the protocol by Bensch et al. (2000), targeting the 5′ region with the primer HaemF. Dye terminator cycle sequencing (BigDye) was conducted on an ABI PRISMTM 3100 capillary sequencer (Applied Biosystems, USA). Sequence editing and alignment were carried out using BioEdit (Hall 1999).

### 2.5 RNA extraction and sequencing

Total RNA was extracted from 20 μL whole blood using 1000 μL TRIzol LS Reagent (Invitrogen, Carlsbad, CA) and homogenized by vortexing from all individuals. Samples were then incubated at room temperature for 5 min before the addition of 200 μl chloroform (Merck KGaA, Darmstadt, Germany). After a further incubation at room temperature for 3 min, the samples were centrifuged at 11,000 rpm for 17 min at 4 °C. The supernatant was then transferred to new tubes and processed using a RNeasy Mini Kit (Qiagen, GmbH, Hilden, Germany). Following the manufacturer’s protocol, 1 volume of 70% ethanol was added to the lysate. The total extracted RNA was shipped on dry ice to Novogene Bioinformatics Technology, Hong Kong, for RNA quality control, DNAse treatment, and rRNA reduction and amplification using the SMARTer Ultra Low Kit (Clontech Laboratories, Inc.). Novogene performed library preparation, cDNA synthesis, and paired-end RNA sequencing using the Illumina HiSeq 2000. We quality checked all demultiplexed RNA-seq reads using FastQC (v.0.10.1) (Andrews 2010).

### 2.6 Bioinformatics analysis

The raw sequence quality was assessed using FASTQC (v.0.10.1) (Andrews 2010). Adapter sequences and low-quality bases were removed using Trimmomatic (v.0.36) (Bolger et al. 2014) with a 4-bp sliding window and a minimum Phred score of 20. High-quality reads were aligned to the canary reference genome (NCBI RefSeq assembly serCan2020) using the splicing-aware aligner STAR (v.2.5.4b) (Dobin et al. 2013). Gene-level quantification was performed using Feature Counts (Liao et al. 2014) to generate a raw read count matrix for downstream statistical analysis.

### 2.7 Differential gene expression level

The read counts for every gene were stored in a file that was statistically analyzed inside the R statistical environment (v.4.5.0) (R Core Team 2023). Read counts were normalized using regularized log transformation to account for potential variation in sequencing depth and the large differences in the number of parasites present in the blood (parasitemia levels). Regularized log transformation of counts was performed to represent the data without any prior knowledge of the sampling design in the PCA and sample distance calculations. The package ggplot2 (Wickham 2016) was employed for the generation of all graphs. Differential gene expression analyses were performed using the DESeq2 package (v.3.21) (Love et al. 2014).

### 2.8 Statistics analyses

Prior to assessing infection-induced transcriptional responses, we evaluated whether the species origin of the inoculated blood matrix elicited host transcriptional modifications independently of malaria infection. To this end, baseline gene expression profiles were directly compared between the two uninfected control groups: birds receiving homologous canary (*Serinus canaria*) blood (Kc) and birds receiving heterologous Eurasian siskin (*Spinus spinus*) blood (Ks). This preliminary analysis allowed us to isolate and quantify the host’s background molecular response associated solely with exposure to a heterologous erythrocyte matrix.

Subsequently, the downstream effects of *Plasmodium* infection were investigated using independent, time-specific cross-sectional factorial designs for each sampling point (8, 12, and 16 dpi). For each independent temporal model, the experimental matrix integrated two factors: (i) Inoculum Source, distinguishing host backgrounds inoculated with canary-derived versus siskin-derived blood matrices, and (ii) Infection Status, segregating infected individuals (Ec, Es) from their respective uninfected controls (Kc, Ks). Factorial differential expression was modeled in DESeq2 utilizing the additive design formula ∼ Inoculum + Infection. By partitioning the variance in this manner, the model mathematically subtracted the transcriptomic signatures attributable to the heterologous blood matrix background prior to estimating the net effect of the parasite infection.

Before the statistical testing at each timepoint, a low-count filter was applied to exclude genes with fewer than 10 cumulative reads across all replicates within that temporal subset, thereby maximizing dispersion estimation accuracy. Statistical significance was determined via Wald tests, and raw p-values were adjusted for multiple testing using the Benjamini–Hochberg False Discovery Rate (FDR) procedure. Genes displaying an FDR 0.05 and an absolute log_2_ Fold Change > 1 were classified as significantly differentially expressed genes (DEGs).

### 2.9 Functional enrichment analyses

To identify biological pathways responding to infection, we performed Gene Ontology (GO) enrichment analysis using g:Profiler (Raudvere et al. 2019). Functional categories were assessed for overrepresentation among upregulated and downregulated DEGs, focusing on biological processes and molecular functions.

To characterize the transcriptomic profile of *P. homocircumflexum*, sequence reads that failed to map to the host canary reference genome during the STAR alignment step were isolated. These host-unmapped raw sequence reads were de novo assembled using Trinity (v.2.15.1) (Grabherr et al. 2011) (please check García-Longoria et al. 2025 for more information about how we proceed with parasite reads). The resulting transcriptome assembly was then structurally and functionally annotated using the Trinotate pipeline (available online: https://github.com/Trinotate/Trinotate.github.io) to identify specific transcript isoforms and genetic features. To assign functional Gene Ontology (GO) terms and molecular functions (such as domain-specific mechanisms or ion transport activities), the curated sequence data was queried against PlasmoDB (Aurrecoechea et al. 2009), an online integrated genomic database for *Plasmodium* parasites. Functional annotations, including GO biological processes, molecular functions, and cell compartment localizations, were retrieved using exact parasite identifier matches to facilitate downstream enrichment analyses.

To evaluate the overall transcriptional investment in host immunity across infection stages, a reference set of avian immune-related genes was compiled through a comprehensive literature search focusing on avian immunology and host-pathogen interactions. A total of 245 curated genes were retained (Supplementary Table S1), comprising 204 innate immunity genes (such as pattern recognition receptors and inflammatory cytokines) and 41 adaptive immunity genes (including components of B-cell and T-cell receptor pathways). Only genes with well-annotated orthologs in the reference genome were included to calculate overall expression intensities (VST normalized counts) per functional category across time point.

## 3. Results

### 3.1. Infection Dynamics

Parasitemia levels were monitored throughout the 32-day experimental period for all groups (Figure 2). Successful infection was confirmed in all individuals from the Ec and Es groups. In both infected cohorts, parasitemia followed a typical trajectory for *P. homocircumflexum*, peaking at 8 dpi. Following the peak, parasitemia levels steadily declined in both groups, reaching near-undetectable levels by 20 dpi (Figure 2). By contrast, individuals in the control groups (Kc and Ks) remained free of parasites for the entire duration of the study.

**Figure 2.**
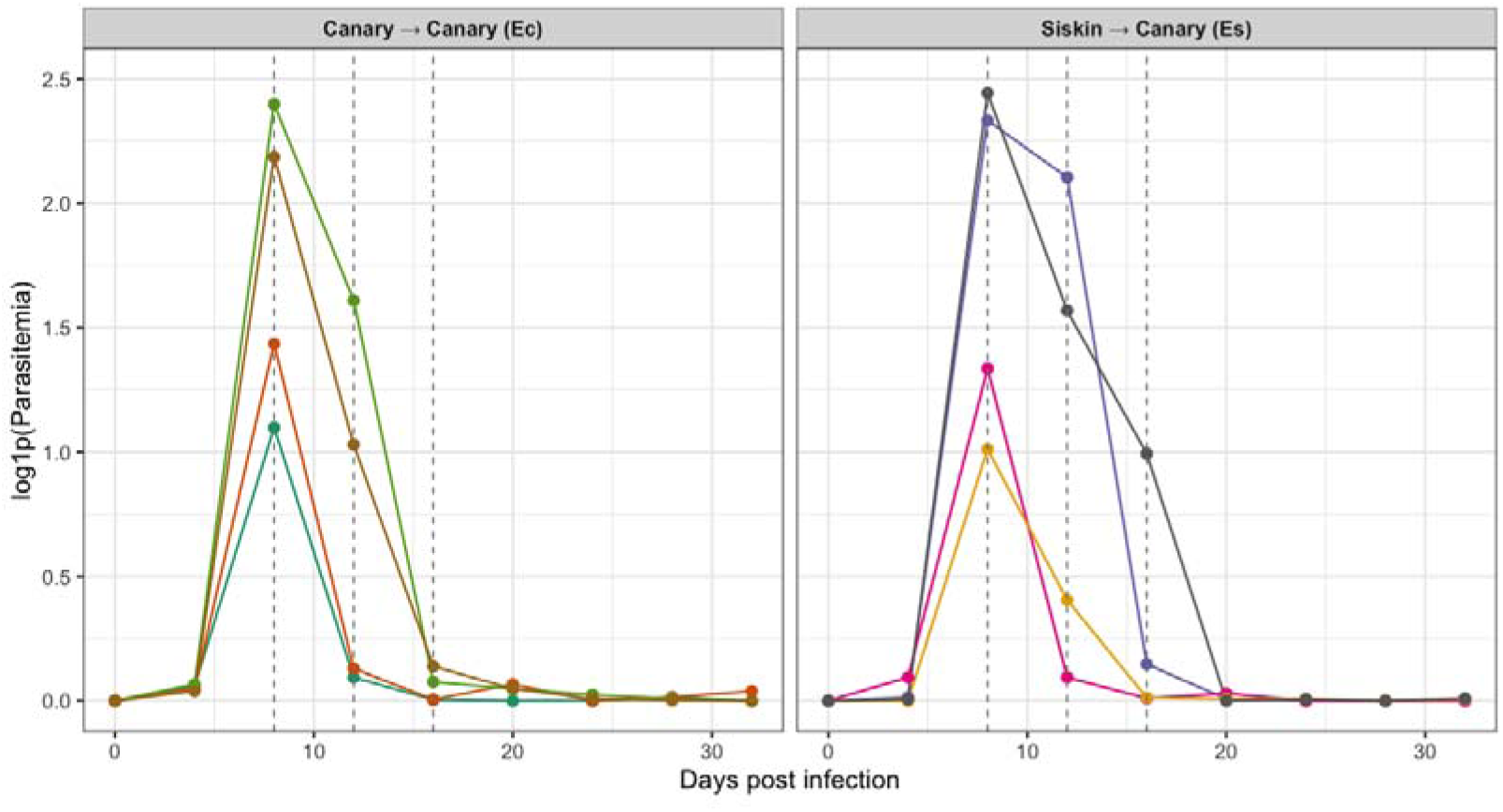
Parasitemia dynamics in canary recipients during the cross-species inoculation experiment. Course of infection over time expressed as log_10_(x+1)-transformed parasitemia (log1p(Parasitemia)) in individual canary (*Serinus canaria*) recipients. The left panel represents the Ec experimental group (Canary Canary), and the right panel represents the Es experimental group (Siskin Canary), both inoculated with *Plasmodium homocircumflexum*. Each coloured line tracks a single individual bird throughout the experiment. Vertical dashed lines indicate the specific sampling points on days 8, 12, and 16 post-infection.

### 3.2. Transcriptional Baseline and Response in Uninfected Controls

To ensure the stability of the experimental system, we first examined the transcriptomic profile of the control canaries at Day 0. Although volcano plots comparing the baseline expression between the Kc and Ks groups show some differentially expressed genes at baseline (Figure 3), functional enrichment analysis indicates that these variations are restricted to normal housekeeping and metabolic processes rather than immune or inflammatory signals (Supplementary Figure S1), confirming that the birds were in a comparable baseline physiological state before inoculations.

**Figure 3.**
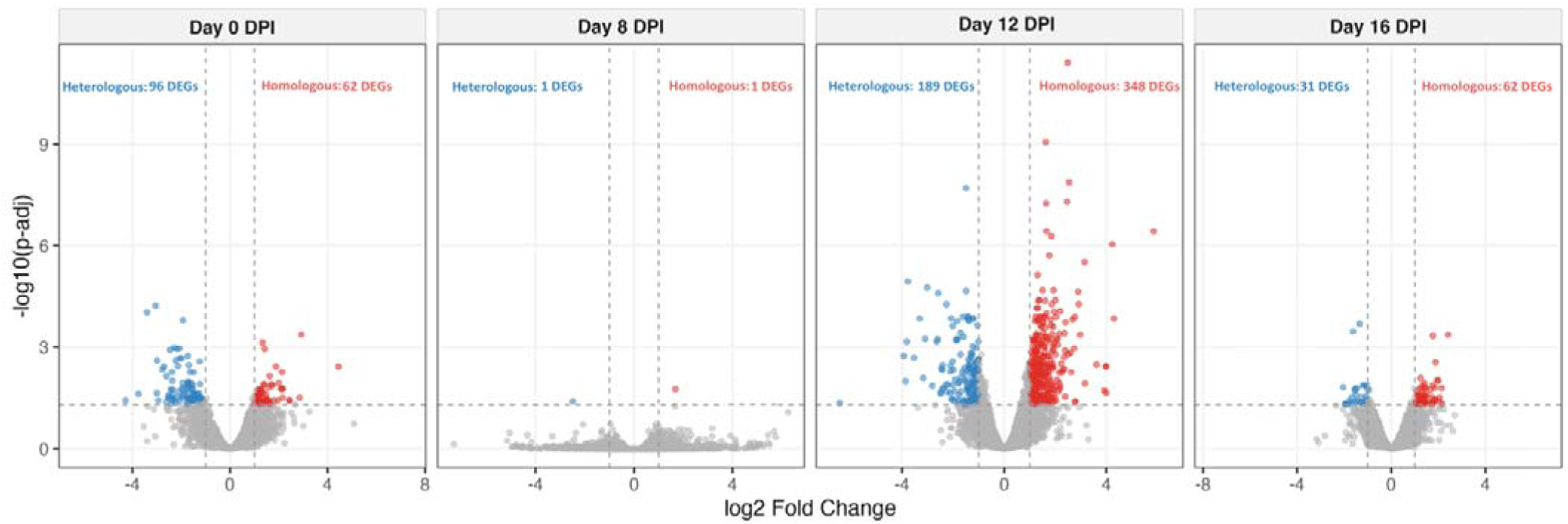
Baseline transcriptional variations and responses between uninfected control groups over time. Volcano plots illustrating differentially expressed genes (DEGs) between the canary canary control group and the siskin canary control group (Kc vs Ks) at distinct sampling points: day 0 (Baseline), day 8, day 12, and day 16 post-infection. Each point represents a single gene plotted by its log_2_Fold Change(x-axis) and statistical significance - log_10_(p-adj) (y-axis). Red dots indicate significantly upregulated genes in the homologous group (log_2_Fold Change> 1, FDR < 0.05), blue dots represent significantly up genes in the heterologous log_2_Fold Change> −1, FDR < 0.05and grey dots show non-significant transcript variations. Dashed horizontal and vertical lines denote the statistical thresholds applied (FDR = 0.05 and log_2_Fold Change=1, respectively). The absolute number of significantly expressed genes (DEGs) is indicated in the top left (heterologous) and top right (homologous) corners of each temporal panel.

The response to clean blood inoculation was evaluated by comparing the Kc (clean canary blood) and Ks (clean siskin blood) groups over time. At 8 dpi, the transcriptional difference was nearly non-existent, with only 2 differentially expressed genes (DEGs) identified (Figure 3). However, at 12 dpi, a significant divergence emerged with 537 DEGs (Figure 3). Functional enrichment analysis at this stage revealed that injecting heterologous blood (Ks) triggered innate immune responses, cytokine-mediated signalling, and macrophage activation (Figure 4). The homologous control group (Kc) remained enriched for homeostatic and metabolic processes, providing a critical baseline to isolate the true host response to malaria (Figure 4).

**Figure 4.**
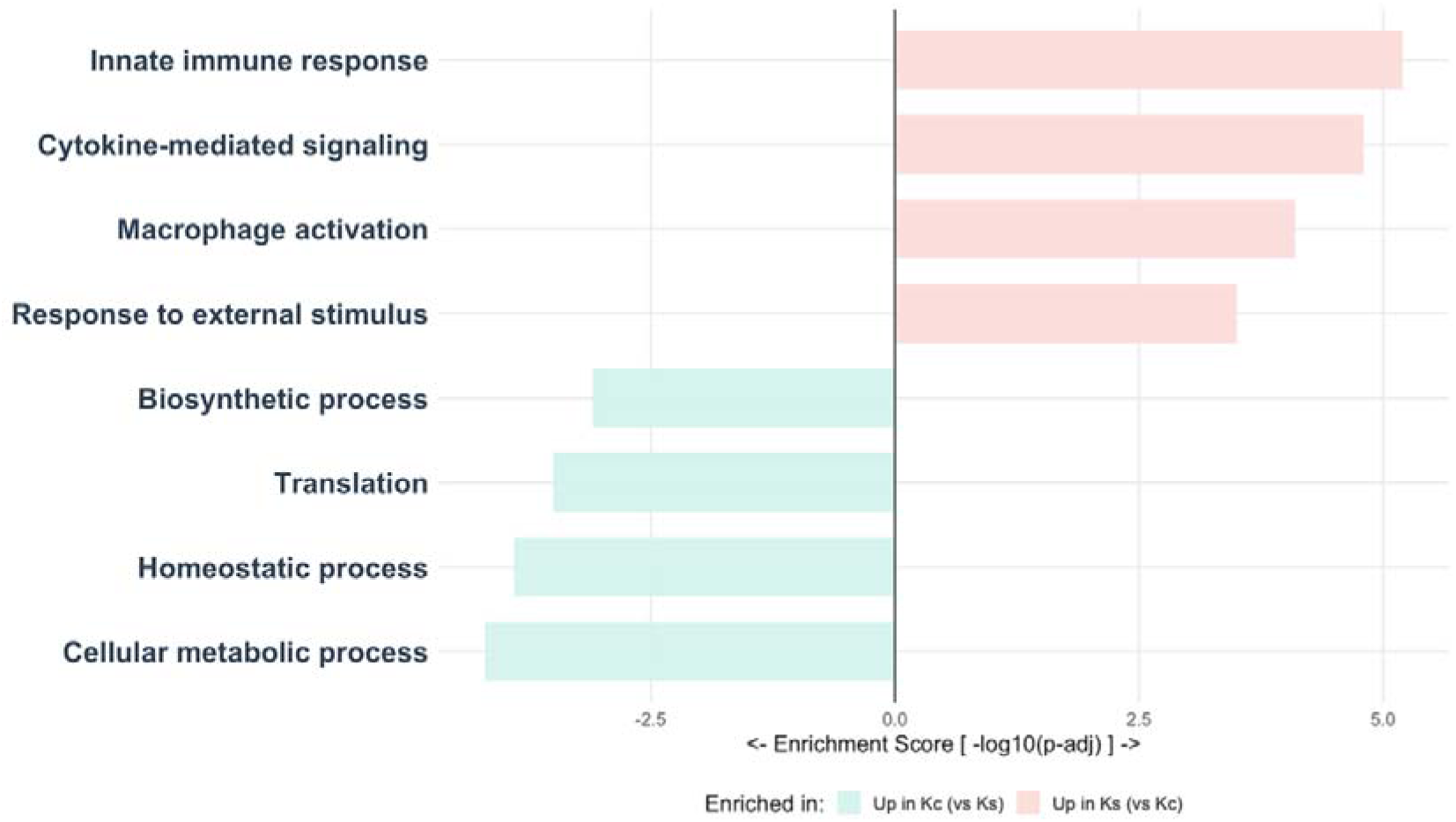
Functional divergence in uninfected host controls at 12 dpi. Divergent Gene Ontology (GO) biological processes enriched between uninfected control groups at day 12 of the experiment. The horizontal barplot displays the enrichment score - for highly significant host pathways. Light green bars (left) represent host biological functions significantly enriched in the canary control group compared to siskins (Up in Kc vs Ks). Light pink bars (right) represent host pathways significantly enriched in the siskin control group compared to canaries (Up in Ks vs Kc).

### 3.3. Corrected Host Transcriptional Response to Malaria

To isolate the specific response to the parasite, we implemented a corrected experimental design that mathematically subtracted the “noise” induced by the heterologous blood matrix (*see Methods section 2.8*). By doing this, it was possible to compare the blood of infected individuals (heterologous vs homologous) whilst eliminating the effect of the blood. Volcano plots using this corrected model showed an intense molecular response during the early and peak stages of infection (Figure 5). At 8 dpi, the host exhibited 2,760 total DEGs (1,614 upregulated and 1,146 downregulated). This response remained robust at 12 dpi with 2,801 total DEGs. By 16 dpi, as the birds effectively suppressed the parasitemia, the number of significant genes dropped sharply to 804 total DEGs (Figure 5).

**Figure 5.**
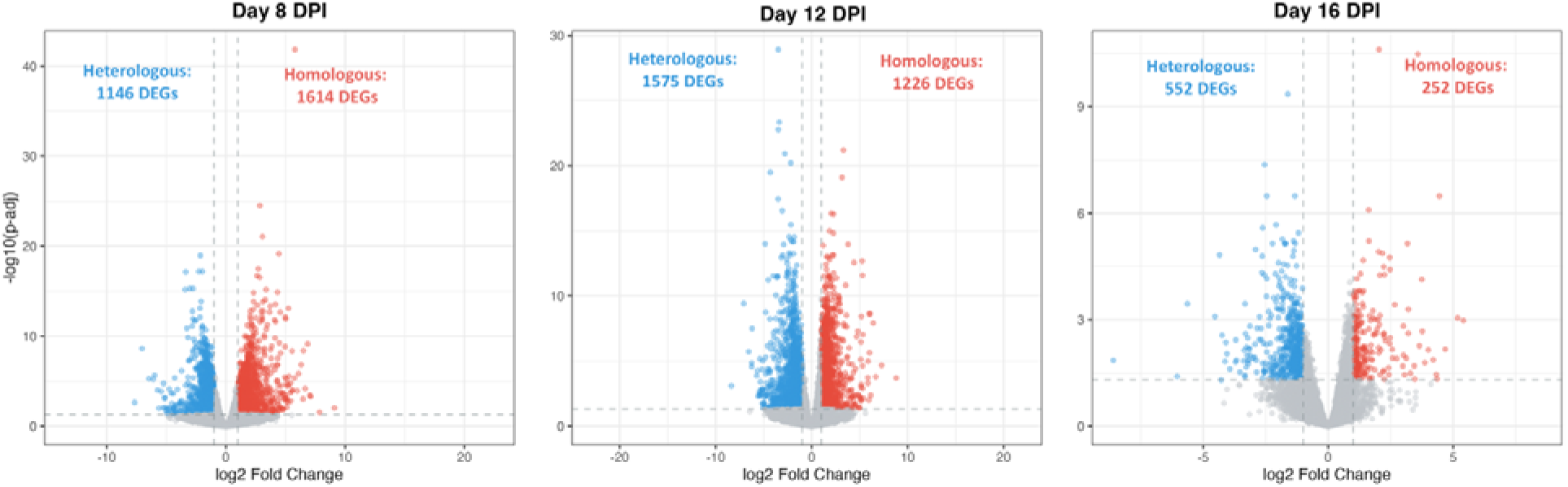
Host transcriptional divergence between heterologous and homologous malaria infections under a corrected design. Volcano plots showcasing the temporal dynamics of differentially expressed host genes (DEGs) at day 8, day 12, and day 16 post-infection. This analysis isolates the net effect of the parasite’s host-shift history by directly comparing the two infection groups after mathematically controlling for and removing the background variance associated with the blood matrix. The x-axis represents the log_2_Fold Change, and the y-axis reflects statistical significance as -log_10_ (p-adjusted). Red dots denote significantly upregulated genes in homologous infections (log_2_Fold Change> 1,FDR < 0.05), blue dots denote significantly upregulated genes in heterologous infections (log_2_ Fold Change < −1, FDR < 0.05), and grey dots indicate non-significant transcripts. Dashed lines mark the established significance thresholds (FDR = 0.05; log_2_Fold Change = 1). The absolute number of significantly altered transcripts is indicated in the top left (heterologous) and top right (homologous) corners of each temporal panel.

### 3.4. Functional Enrichment and Temporal Evolution

Functional enrichment analysis revealed highly divergent host response patterns at 8 dpi between the experimental cohorts (Figure 6). At 8 dpi, the Ec group exhibited a profound overrepresentation of terms related to catalytic activity, whereas the Es group uniquely mobilized structural ribosomal constituents and RNA-binding mechanisms (Figure 6). Notably, these stark initial discrepancies largely disappeared by 12 and 16 dpi, giving way to more synchronized or attenuated profiles (Figure 6). This early divergence is strongly supported by the selective upregulation of core regulatory hub genes specific to each cohort (Figure 7), including major signalling and homeostatic drivers (AKT1, CDK6, DICER1, GSK3B).

**Figure 6.**
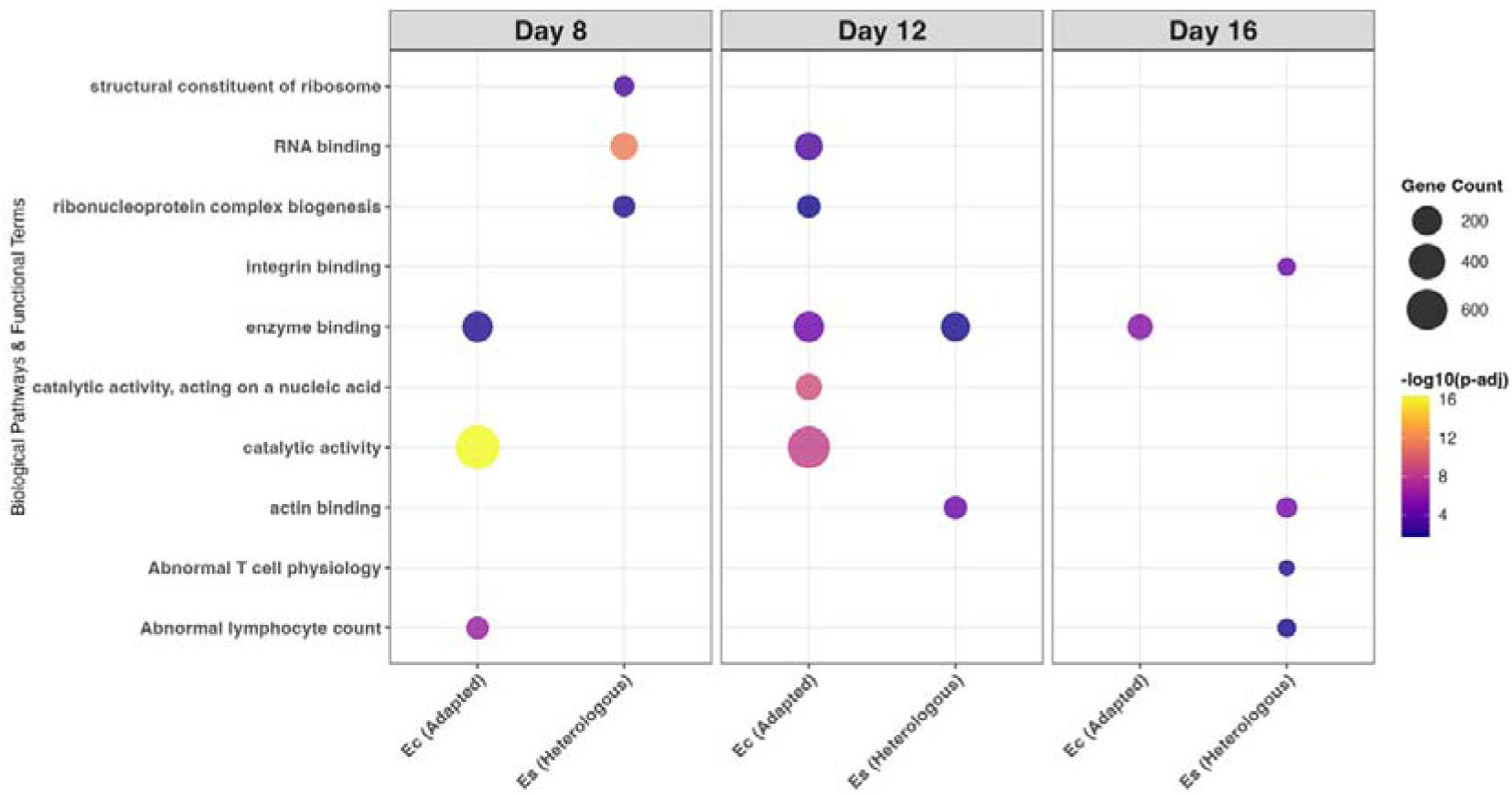
Functional enrichment evolution of the host response to malaria. Comparison of enriched biological pathways and functional terms between adapted (Ec) and heterologous (Es) experimental infection groups across three post-infection sampling points: Day 8, Day 12, and Day 16. Enriched functional terms are plotted on the y-axis, grouped by temporal panels. The dot size represents the “Gene Count” (the total number of differentially expressed genes associated with each specific term), while the dot colour indicates the statistical significance level, expressed as -log_10_(p-adjusted), ranging from low significance (dark blue/purple) to highly significant enrichment (yellow).

**Figure 7.**
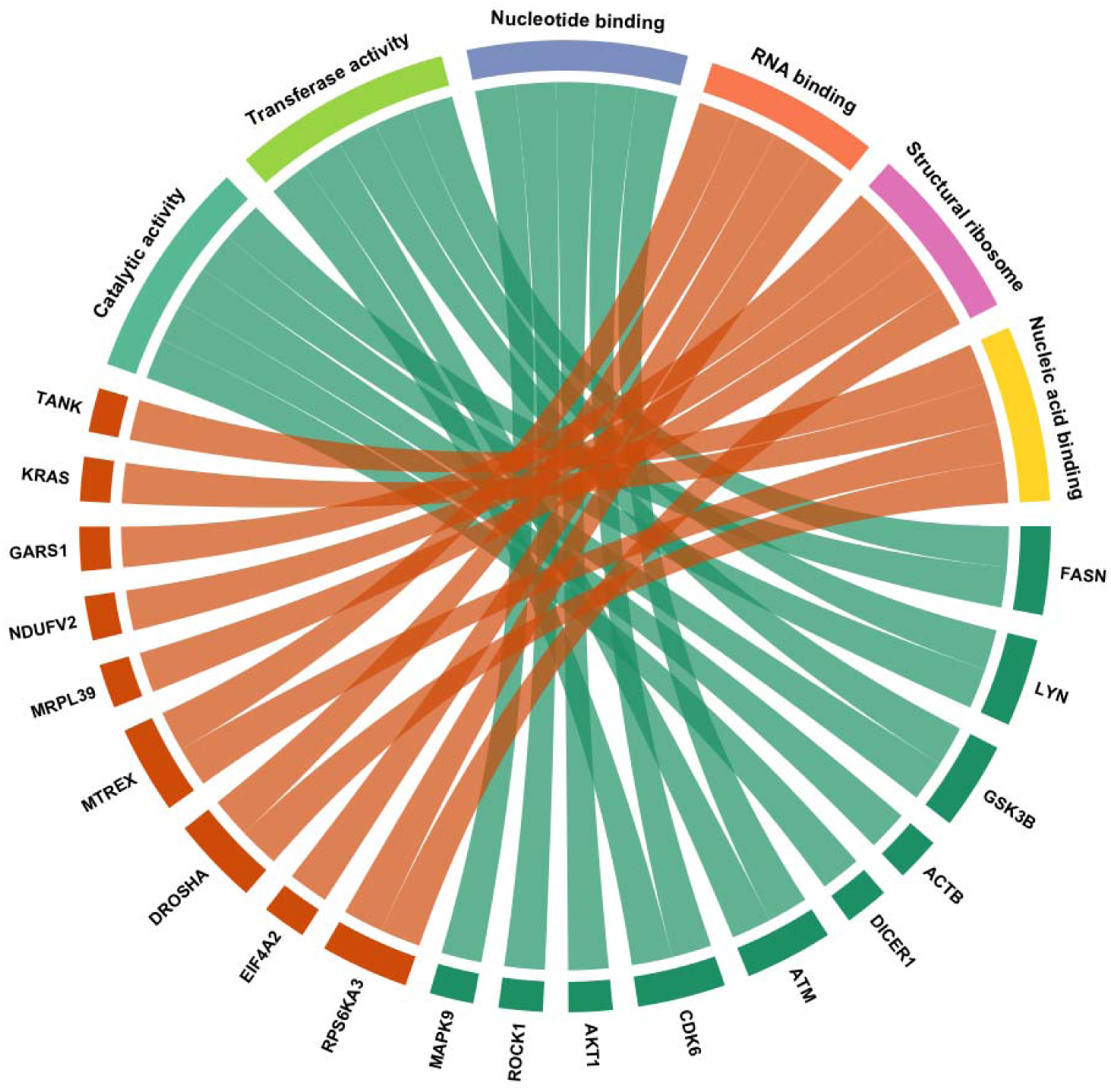
Chord diagram illustrating the relationship between key differentially expressed host genes and enriched functional categories at 8 dpi. Circle plot mapping representative host genes (bottom and left segments, orange and teal blocks) to their corresponding Gene Ontology (GO) functional terms and molecular activities (top and right segments, multi-coloured blocks). The green (teal) blocks identify master regulator host genes exclusive to the adapted infection (Ec), whilst the orange blocks indicate host genes specific to the heterologous infection (Es), highlighting the absence of genes shared between both pathways at this critical point in the infection. Coloured ribbons connect individual host genes to the specific functional categories in which they are involved. The width of the ribbons is proportional to the gene’s involvement across categories, highlighting core regulatory hubs (e.g., kinases like *AKT1*, *CDK6*, *GSK3B*, and *ATM*) linking catalytic, transferase, and nucleotide/RNA-binding processes.

### 3.5. Transcriptional Dynamics of Parasite and Host Immune Response

At 8 dpi, *P. homocircumflexum* exhibited high transcriptional activity in the Ec and Es cohorts (Figure 8). Isoform quantification revealed enriched transcripts in key functional categories, most notably Metal ion transport (ZIP Zinc) with over 250 transcripts, alongside Intracellular signalling (PI3/PI4) and Protein ubiquitination (RING) (Figure 8). By 12 dpi, a stark transcriptomic divergence occurred between parasites that was transferred from different donors: while the parasite population in the Ec group maintained elevated levels of enriched isoforms for Metal ion transport, other categories such as Kinases showed comparable levels between cohorts, whereas the Es group showed a significant reduction in other targeted pathways (Figure 8).

**Figure 8.**
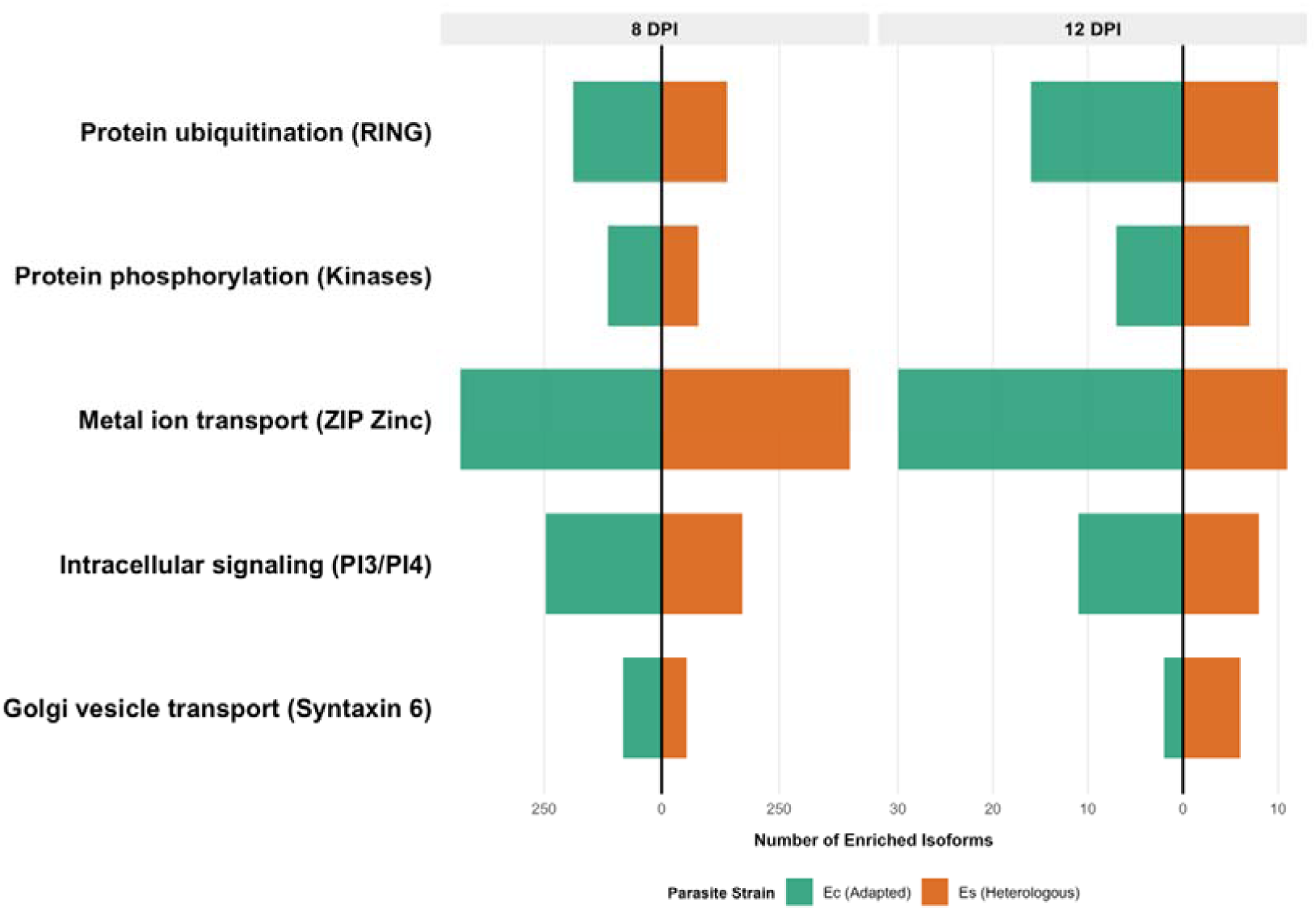
Comparative transcriptomic response of *Plasmodium homocircumflexum* over time. Mirror barplot showing the direct quantification of enriched parasite isoforms across key biological categories at 12 days post-infection (12 dpi, left panel) and 8 days post-infection (8 dpi, right panel). Green bars represent the adapted parasite population (Ec, green), and orange bars represent the heterologous parasite population (Es, orange). The x-axis indicates the absolute number of enriched isoforms, noting the axis scale divergence between early widespread expression at 8 dpi (counts exceeding 250 transcripts) and the targeted transcriptional divergence at 12 dpi.

To determine if these specific regulatory divergences translated into broader shifts in immune activation, we analysed overall expression intensities for adaptive and innate immunity (Supplementary Figure S2). Despite the functional divergence observed in specific pathways (where Ec and Es groups diverge in which functional networks they mobilize, as shown in Figures 6–8), the overall magnitude of immune-gene activation remained relatively consistent between the groups across the time course. Both adaptive and innate expression profiles showed comparable global investment between parasite populations, with no significant differences overall, except for a slight, statistically significant variation observed at 12 dpi in the innate arm (Supplementary Figure S2).

## Discussion

Despite the widespread use of avian models in evolutionary biology and ecology, the systemic and molecular impacts of heterologous blood transfusions between bird species have historically remained understudied (Degernes et al., 1999; Martinho et al. 2009). Although conventional paradigms assumed that birds – lacking conventional mammalian blood typing systems – experience minimal physiological friction during transfusions (Degernes et al., 1999; Cital et al. 2016), recent evidence challenges this view. Notably, the discovery of an erythrocyte glycan system in birds, analogous to the human P1PK blood group, suggests a robust molecular mechanism for immune recognition during interspecies transfers (Bereznicka et al., 2026). Here, we resolve the functional and transcriptomic landscapes altered by blood transfusions between species. Given that the domestic canary (*Serinus canaria*) and the Eurasian siskin (*Spinus spinus*) are closely related phylogenetically, a muted physiological response might be reasonably anticipated. Crucially, our results refute this assumption: canaries receiving parasite-free heterologous blood mounted a massive, systemic immune response characterized by 537 differentially expressed genes (DEGs) at 12 dpi (Figure 4), heavily driven by cytokine signalling and macrophage activation. This profound host reactivity reveals a major confounding variable that has been traditionally overlooked in experimental infection studies utilizing blood-borne inoculum. Ultimately, to avoid systemic misinterpretation of host RNA dynamics (Lee et al., 2018), our findings demonstrate that future avian immunological and expression research must rigorously account for the baseline immune signatures induced by the transfusion matrix itself.

This confounding effect is particularly critical for historical avian malaria frameworks, where blood-stage parasites are routinely inoculated via heterologous donors without adjusting for the transcriptomic noise of the donor’s erythrocyte matrix (Videvall et al. 2020; Paxton et al. 2023). By applying an independent cross-sectional factorial design at each distinct sampling window, we successfully partitioned out the background transcriptomic variance of the blood matrix, enabling us to isolate the clean, parasite-specific signal. When we eliminate the baseline effect of the blood and focus strictly on the net effects of the parasite, we find significant transcriptional differences between homologous and heterologous as early as 8 dpi. Importantly, the fact that the uninfected heterologous matrix continues to drive an active independent profile at 12 dpi underscores the persistence and non-linear nature of this background noise, validating our independent temporal correction and preventing chronological misinterpretation of host reactivity. This divergence in the response of the bird is not a subtle nuance, but a profound discrepancy in the host’s cellular priorities.

In the homologous group (adapted infection), we observed a massive catalytic overload from 8 dpi onwards, characterised by frenetic enzymatic activity and transcriptomic signatures indicative of altered lymphocyte dynamics and mobilization, rather than simple changes in systemic cell counts, by deploying an overrepresentation of pathways associated with a catalytic overload mechanism regulated by critical kinase hubs (*AKT1*, *CDK6*, *GSK3B*, and *ATM*). This functional profile might suggest that, as a parasite already adapted to the canary’s environment, *P. homocircumflexum* could be more effective and aggressive at invading the tissues of the host. Consequently, the canary enters a defensive emergency phase characterized by the enrichment of pathways driving mitotic cell cycle disruption and cellular stress signaling, reflecting the high physiological cost and systemic inflammation of an unchecked adapted proliferation (Ellis et al. 2015; Sheppard et al. 2024). In contrast, the response to heterologous infection is marked by the prioritization of ribosome biogenesis, binding to RNA and, subsequently, structural integrity via integrins and actin. This ability to implement structural regulation rather than an energetic metabolic panic response suggests that the bird shifts its transcriptome toward a more efficient early management of the infection. This immunological advantage for the host, however, could not be a choice made by the bird itself, but rather the direct result of a window of opportunity opened by the pathogen’s internal state and its cell-cycle coordination with the host. The key to this divergence could lie in the fact that the heterologous parasite, hampered by its previous history, fails to synchronise its metabolic and replication cycles at the same rate as the adapted parasite population – a phenomenon of adaptive lag described in other parasite species.

This divergence observed at 8 dpi in the RNA expression profile of the host could correspond to the differences in RNA expression recently observed in the parasite (García-Longoria et al. 2025). In this regard, García-Longoria et al. (2025) suggested that parasite expression depended on the origin of the infection via epigenetic memory, whilst after that time window the differences depended on the environment (host) rather than the parasite population. In support of this idea, we have found that in the homologous group, the parasite overexpresses zinc transporters, kinases and RING ubiquitin ligases (Figure 8), which could correlate with the bird’s functional profile. In contrast, the adaptive delay in the heterologous group limits this initial response, allowing the bird to implement RNA expression based on structural integrity (actin and integrins) rather than an emergency response. This real-time coordination demonstrates that pathogenicity is not a fixed trait, but rather an emergent outcome of the parasite’s previous history, which dictates the host’s molecular fate from the very earliest stage of infection

In conclusion, our study demonstrates that the pathogenicity of a generalist parasite should not be understood as a static trait of its genome, but rather as an emergent property determined by its recent history of host switching (Prati et al. 2022). By establishing, for the first time, a filtered avian malaria infection model that mathematically isolates immunogenic noise from the blood matrix (Frank and Schmid-Hempel 2008; Gandon 2018; Mahanta et al. 2018), we have revealed that the pathogen’s adaptive lag acts as a critical determinant of virulence in nature. This phenomenon suggests that the severity of infectious outbreaks in complex ecosystems depends intrinsically on host-switching pathways (Agosta et al. 2010; Handel et al. 2010; Gupta et al. 2020) and on the parasite’s epigenetic flexibility to navigate divergent metabolic landscapes (García-Longoria et al. 2025). By integrating the molecular trajectories of both organisms into a coordinated dialogue, this work validates the need to adopt a systems-based paradigm for investigating infectious diseases, in which the host and pathogen are no longer viewed as isolated entities but are treated as dynamic partners (Lee et al. 2018; Mukherjee et al. 2021). The future challenge for evolutionary biology will lie in deciphering the specific epigenetic code – methylation marks and histone modifications – that underpins this transcriptional memory, ultimately enabling us to predict and mitigate the impact of pathogens that conquer new ecological frontiers.

## Funding

LGL was funded by Junta de Extremadura (IB24107). VP and AJ were funded by a grant (No. S-MIP-25-42) from the Research Council of Lithuania. O.H. obtained funding from the Swedish Research Council (VR 2016-03419 and 2021-03663).

---

**Figure S1.**
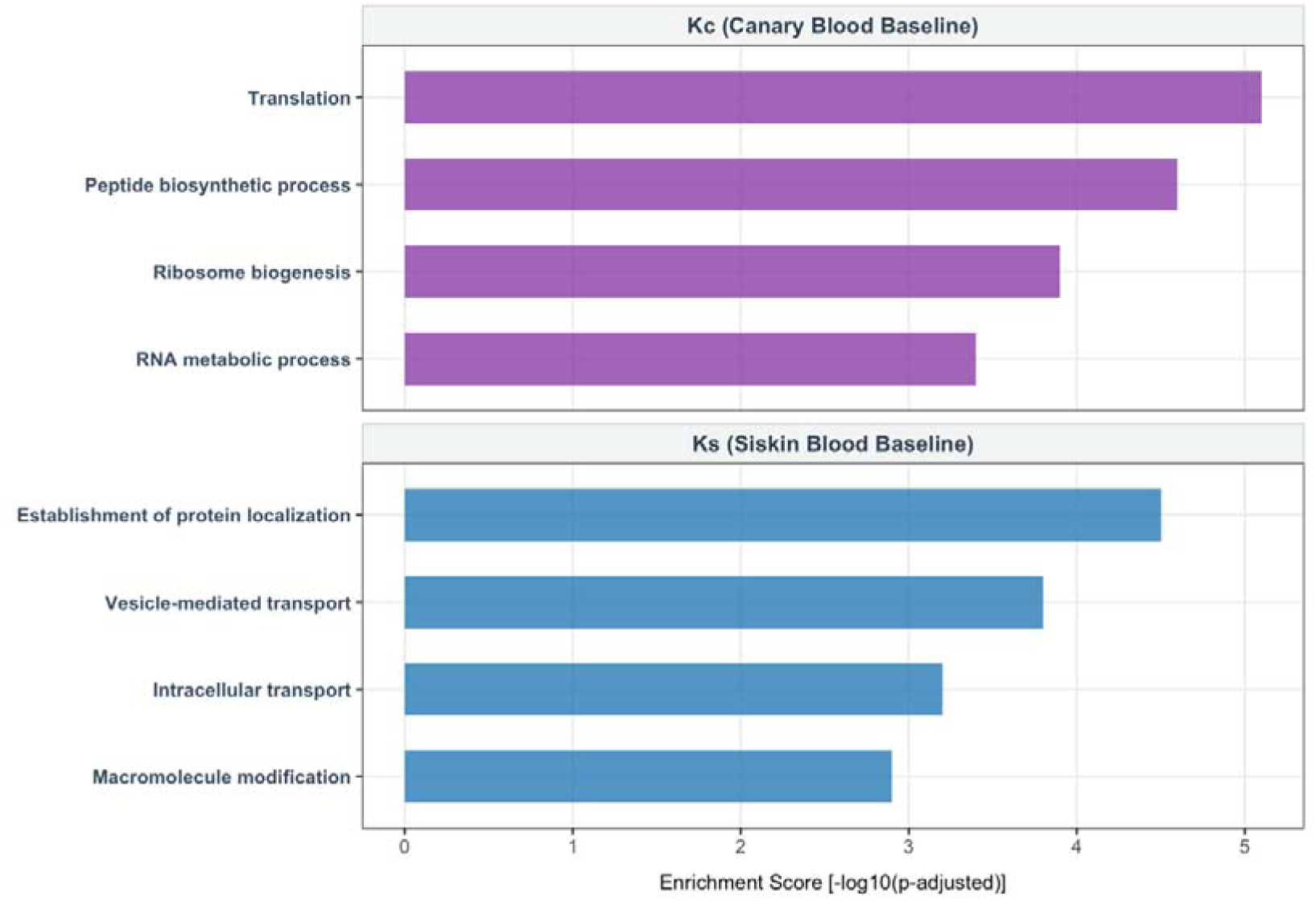
Functional enrichment analysis of baseline metabolic variations at day 0 (pre-infection). Significantly enriched biological processes in healthy control groups before experimental inoculation. The top panel illustrates the baseline profile for the canary control group (Kc, purple bars), showing metabolic fluctuations associated with standard housekeeping processes (e.g., translation, ribosome biogenesis). The bottom panel illustrates the baseline profile for the siskin control group (Ks, blue bars), highlighting pathways related to cellular transport and localization. Bars represent the enrichment score calculated as -log_10_(p-adjusted).

**Figure S2.**
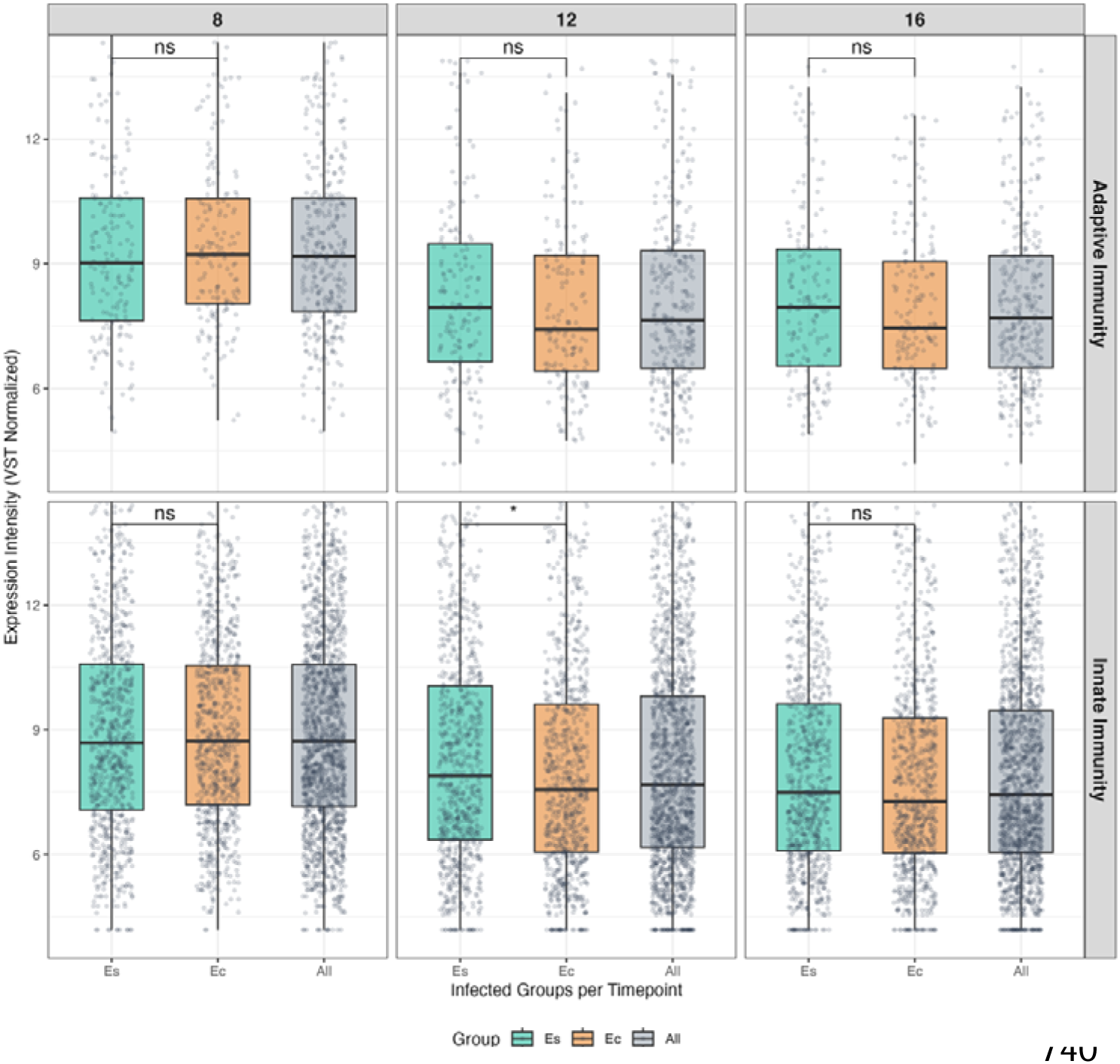
Temporal dynamics of avian immune system profiles. Pairwise statistical comparison of host gene expression intensities across infection groups and experimental timepoints. Boxplots represent Variance Stabilizing Transformed (VST Normalized) expression data for genes categorized under Adaptive Immunity (top panels) and Innate Immunity (bottom panels) at days 8, 12, and 16 post-infection. Experimental cohorts are color-coded as follows: heterologous infection (Es, green), adapted infection (Ec, orange), and the combined global profile (All, grey). Individual data points are overlaid as a transparent jitter plot to display real gene dispersion. Pairwise statistical comparisons between Es and Ec are indicated by brackets at the top of each panel (*: p < 0.05, ns: non-significant).

## Notes

https://www.ncbi.nlm.nih.gov/sra/PRJNA1518241

### Competing Interest Statement

The authors have declared no competing interest.

